# Harnessing defective interfering particles and lipid nanoparticles for effective delivery of an anti-dengue virus RNA therapy

**DOI:** 10.1101/2024.04.23.590664

**Authors:** Min-Hsuan Lin, Pramila Maniam, Dongsheng Li, Bing Tang, Cameron Bishop, Andreas Suhrbier, Lucy Wales-Earl, Yaman Tayyar, Nigel A. J. McMillan, Li Li, David Harrich

**Affiliations:** Program of Infection and Inflammation, QIMR Berghofer Medical Research Institute, Herston, Queensland 4006, Australia; Global Virus Network (GVN) Center of Excellence, Australian Infectious Disease Research Centre, Brisbane, QLD, Australia; Menzies Health Institute Queensland and School of Pharmacy and Medical Science, Griffith University, Gold Coast, Australia; Prorenata Biotech, Molendinar, Queensland, Australia; Australian Institute for Bioengineering and Nanotechnology, the University of Queensland, St Lucia, Queensland 4072, Australia

**Keywords:** defective, interfering, RNA, DIPs, LNPs, dengue, therapeutic, antiviral, mouse

## Abstract

Presently, no approved antiviral drug targets dengue virus (DENV) infection. Treatment mainly relies on supportive measures, while DENV vaccines’ efficacy varies based on factors like vaccine type, circulating DENV serotypes and vaccinated population. This study explores using defective interfering particles (DIPs) and lipid nanoparticles (LNPs) to deliver an anti-DENV defective interfering RNA, known as DI290. Results showed that both DIPs and DI290 loaded LNPs (LNP-290) effectively suppressed DENV infection in human primary monocyte-derived macrophages (MDMs), THP-1 macrophages and human fibroblasts, representing cell types naturally targeted by DENV. Furthermore, LNP-290 demonstrated >log_10_ inhibition of DENV viral loads in IFNAR-deficient mice, which lack functional type I interferon (IFN) receptors. DI290-mediated inhibition was also effective in IFN regulatory factor 3 and 7 double knockout mice. RNA-Seq data from LNP-treated C57BL/6J mice, IFNAR-deficient mice and human MDMs treated with LNPs or DENV DIPs illustrated DI290 treatment heightened IFN responses, particularly IFNγ, as well as IFNα/β and IFNλ. DI290 thus induces a broad range of IFN responses, with IFNγ and IFNλ providing anti-viral activity when IFNα/β responses are absent. Mice administered LNP-290 also did not manifest acute overt clinical signs. In summary, these experiments suggest DI290’s potential as a therapeutic approach for combating DENV infection.

## Introduction

DENV is a single-stranded RNA virus from the *Flaviviridae* family transmitted to humans by the *Aedes aegypti* mosquito. It is estimated that approximately 390 million people are infected with dengue virus each year, making it one of the most common mosquito-borne viral diseases worldwide.^1^ The disease can manifest with a wide range of symptoms, from mild flu-like illness to severe and potentially fatal forms such as dengue hemorrhagic fever (DHF) and dengue shock syndrome (DSS). The development of antiviral drugs for dengue virus is crucial as they can help reduce the severity of the infection and prevent complications. Traditional antiviral drugs typically function by directly targeting the virus, either by impeding its replication or by obstructing its entry into and infection of host cells. Nonetheless, because of the absence of approved specific antiviral treatments and restricted availability of vaccines for dengue virus,^2,3^ virus management primarily hinges on mosquito control measures, prompt detection and treatment of cases, and prevention of complications like fever, dehydration, bleeding and shock.

We described DI290 RNA, a short 290-nucleotide defective interfering (DI) RNA derived from DENV.^4–7^ DI RNAs are produced by RNA viruses due to errors in the viral RNA replication process. These errors lead to incomplete or shortened copies of the viral genome, which lack some or all of the genes necessary for viral replication. ^reviewed^ ^in^ ^8,9^ DI290 RNA retains the intact 5’ and 3’ untranslated regions (UTRs) of the viral genome, but it lacks all open reading frames. DI RNAs, including DI290 RNA, can disrupt the replication of the parent virus through various mechanisms.^8,10^ Firstly, DI RNAs are suggested to compete with the parent virus for limited resources within the host cell, such as enzymes and nucleotides essential for viral replication. Secondly, DI RNAs may hinder the replication of the parent virus by acting as a template for RNA synthesis. The viral polymerase replicates DI RNAs, producing more DI RNAs rather than viral genomic RNA. Previously, we demonstrated that DI290 RNA replicated in cells infected with DENV or Vero E6 cells expressing the DENV RNA polymerase ^5^. Thirdly, DI RNAs might hinder the assembly and release of viral particles. They can compete with the viral genome for packaging signals crucial for viral assembly, leading to the production of defective viral particles, known as DIPs, which cannot infect new cells. In summary, DI RNAs contribute to controlling virus replication by restricting available resources, competing for RNA synthesis and disrupting viral assembly and release. Through these mechanisms, DI RNAs diminish the virus’s capacity to propagate and induce disease.

We employed stable cell lines to synthetically generate DENV-based DIPs, which were purified and concentrated ^6^. Our findings demonstrated that DIPs exhibit broad-spectrum antiviral activity by eliciting innate immune responses in cells.^6^ There is evidence suggesting that DI RNAs can bind the RLRs, RIG-I and melanoma differentiation-associated protein 5 (MDA5), activating this signaling pathway and initiating antiviral responses in host cells.^11,12^ A direct interaction between the DENV 5’ UTR and RIG-I has been reported.^13^ Upon recognition of a target RNA, RLRs undergo a conformational change, exposing its caspase activation and recruitment domains (CARDs). These exposed CARDs of RLRs then interact with the mitochondrial antiviral signaling protein (MAVS) located on the outer mitochondrial membrane. This interaction initiates a signaling cascade involving the activation of various downstream molecules, including TBK1 (TANK-binding kinase 1) and IKKε (I-kappa-B kinase epsilon). MAVS deficiency in cells abrogates the ability of DI RNA to induce antiviral responses to DI RNAs.^14^ TBK1 and IKKε subsequently phosphorylate IFN regulatory factor (IRF)3, IRF7 and NF-κB (Nuclear Factor-kappa B), respectively. Phosphorylated IRF3, IRF7 and NF-κB translocate to the nucleus and induce the transcription of type I/III IFNs and other proinflammatory cytokines reviewed in ^15^. In general, DI RNAs originating from various RNA viruses have been observed to activate innate immunity and prompt an antiviral response in host cells ^14,16–19^. This event aids in regulating viral replication and dissemination.

In this study, we explored DI290’s capacity to impede DENV replication in primary human macrophages, foreskin-derived fibroblasts and in two mouse models of DENV-2 infection. Our findings indicate that DI290 shows promise as an anti-DENV agent by activating IFN responses, capable of diminishing virus replication and decreasing viral titers *in vivo*. Consequently, it holds potential as a therapeutic intervention against DENV infection.

## Results

### DI290 RNA protects human macrophage and fibroblasts from dengue virus infection

Transmission of DENV by mosquitoes through skin bites involves several dermal and epidermal cells that are susceptible to infection, including keratinocytes, dendritic cells, Langerhans cells, and fibroblast cells.^20–22^ After entering the bloodstream, DENV predominantly infects monocytes and macrophages, acting as a cellular reservoir for virus replication.^23–27^ In this study, our first objective was to examine whether DENV DIPs induce interferon responses in human primary MDMs, similar to observations made previously in Huh7 hepatoma cells. ^6^ MDMs treated with DENV DIPs exhibited a transient and rapid increase in IFN-β mRNA levels at 2 hours post-treatment, alongside transiently elevated levels of IFN-λ mRNA at 24 hours (Fig. 1A-B, respectively). Moreover, RIG-I showed significant upregulation at 2 hours post-treatment (Fig. 1C), while both RIG-I and interferon-stimulated gene 15 (ISG15) were upregulated at subsequent time points (Fig. 1C and Fig. 1D, respectively). Similar observations were made in THP-1 macrophage cells treated with DIPs, demonstrating notable upregulation of RIG-I and ISG15 (Supplementary Fig. 1A-B). Additionally, HFF-1 cells, a human fibroblast cell lines, exhibited approximately a 2-fold increase in IFN-β mRNA levels and a 20-to 40-fold increase in RIG-I and ISG15 mRNA levels, respectively, at 24 hours post-DIP treatment (Fig. 2A-D).

**Figure 1.**
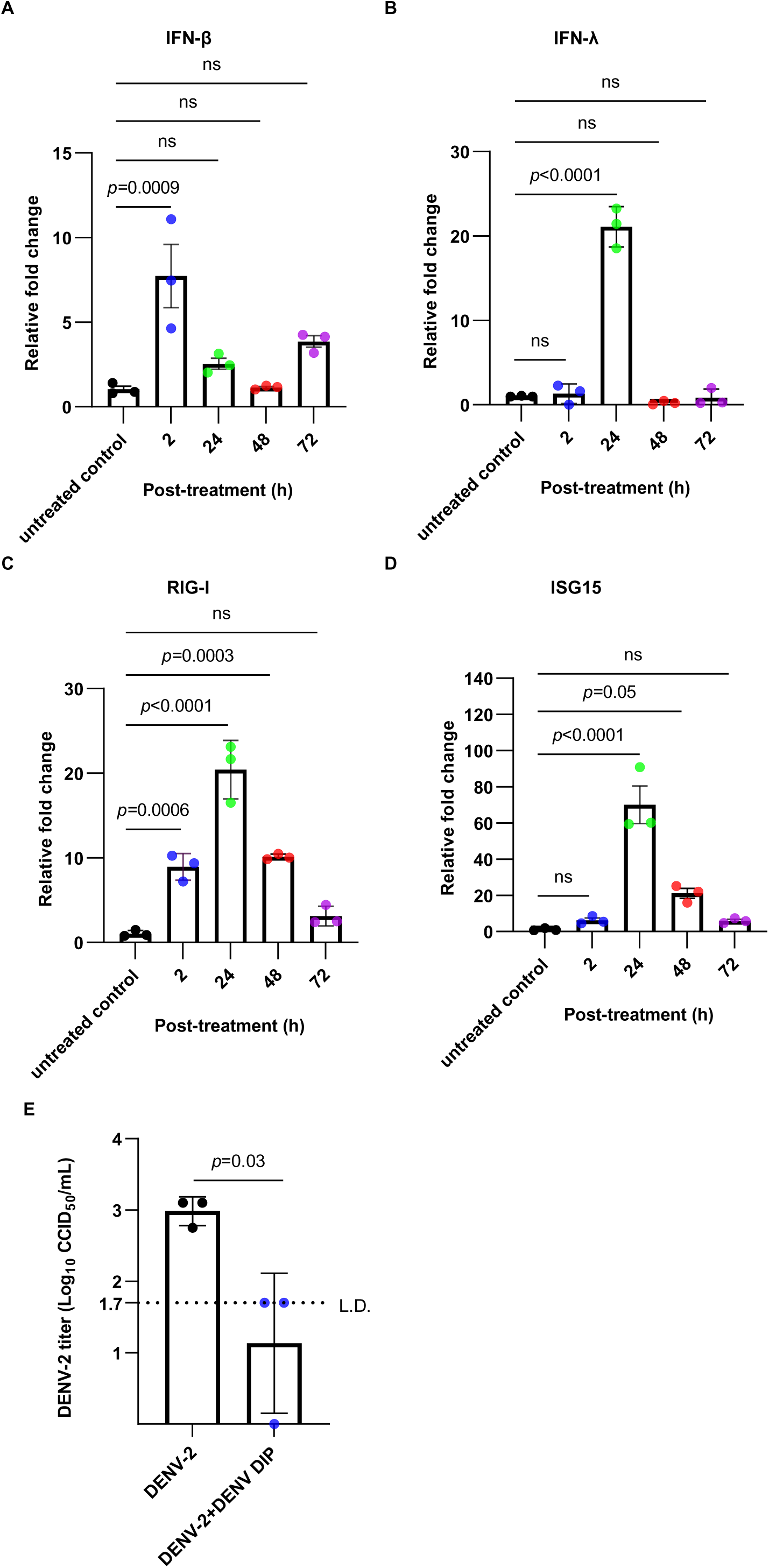
DENV DIPs stimulate host innate immune responses in human primary MDMs and inhibit DENV-2 replication. (A-D) Monocytes were isolated from human PBMCs using CD14 positive selection and maturated to MDMs with M-CSF and GM-CSF for 5 days. MDMs were then treated with DENV DIPs (at a dosage equivalent to 1,000 DI290 RNA copies (=0.2 fg) per cell) for 2, 24, 48 and 72 h. Total RNA was extracted from the cells and the levels of IFN-β, IFN-λ, RIG-I and ISG15 mRNA were quantified by RT-qPCR. The fold change relative to the untreated control cells was calculated (n=3). (E) MDMs were infected with DENV-2 (MOI=1 CCID_50_ per cell). After 3 h, the cells were washed with 1X PBS and incubated with culture medium containing DENV DIP (at a dosage equivalent to 1,000 DI290 RNA copies (=0.2 fg) per cell). DENV-2 titers in culture supernatant were measured by CCID_50_ assay at 3 days post-infection (n=3). The data are shown as the mean ± standard of deviation (SD). Statistical analysis was performed by one-way ANOVA (A to D) or student’s *t*-test (E). L.D.: limit of detection. ns: not significant.

**Figure 2.**
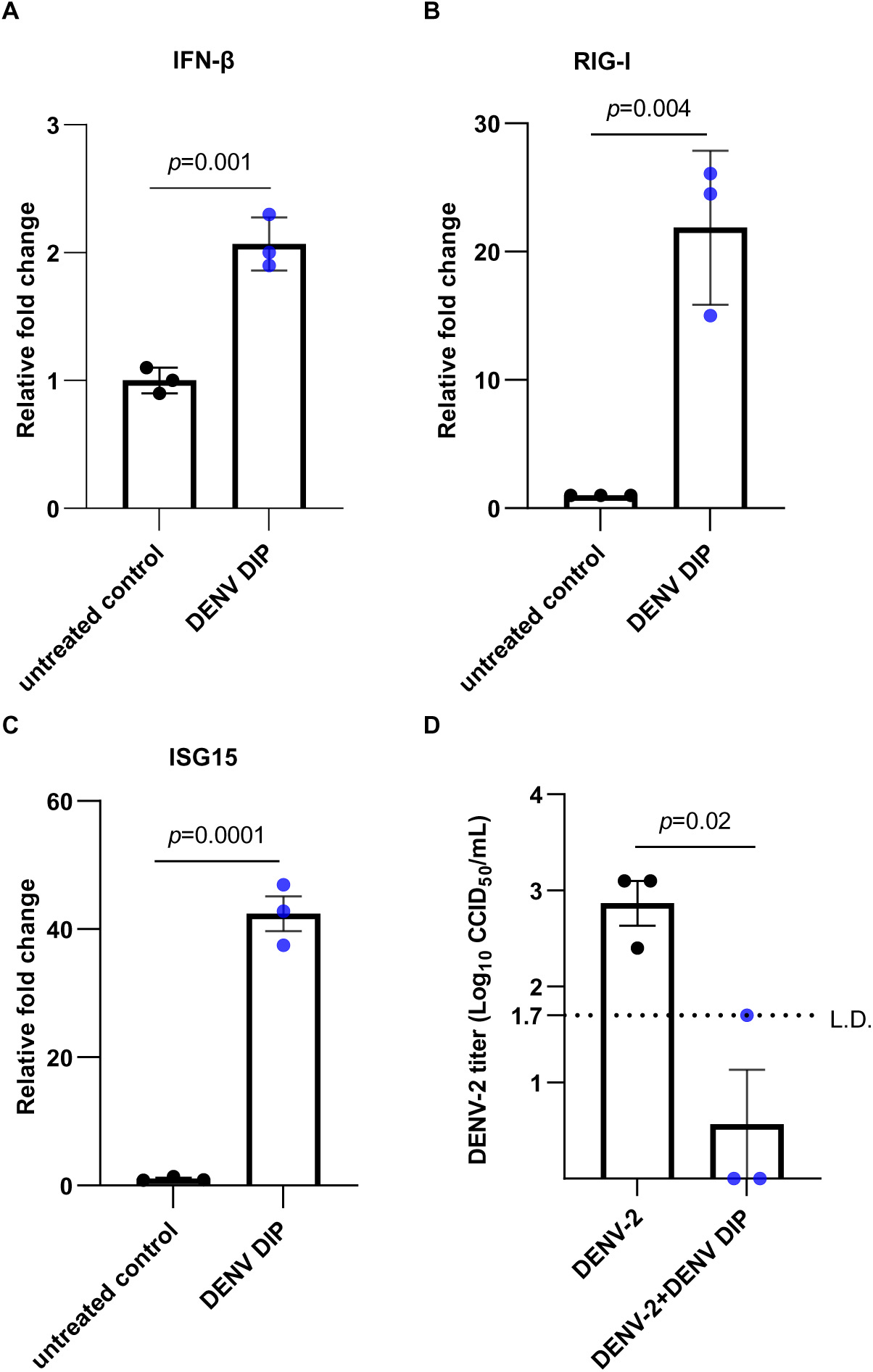
DENV DIPs activate IFN responses and inhibits DENV-2 in HFF-1 human fibroblast cells. (A-C) HFF-1 cells were treated with DENV DIP (at a dosage equivalent to 1,000 DI290 RNA copies (=0.2 fg) per cell) for 24 h. Total RNA was extracted from the cells and the levels of IFN-β, RIG-I, and ISG15 mRNA were quantified by RT-qPCR. The fold change relative to the untreated control cells was calculated (n=3). (D) HFF-1 cells were infected with DENV-2 (MOI=0.1 CCID_50_/cell). After 3 h, the cells were washed with 1X PBS and incubated with culture medium containing DENV DIP (at a dosage equivalent to 1,000 DI290 RNA copies (=0.2 fg) per cell). DENV-2 titers in culture supernatant were measured by CCID_50_ assay at 3 days post-infection (n=3). Data are shown as the mean ± SD. Statistical analysis was performed by student’s *t*-test.

Subsequently, all three cell types were infected with DENV-2 for 3 hours, employing a MOI of 0.1 for HFF-1 cells and a MOI of 1.0 for MDMs and THP-1. After removing the DENV inoculum, the cells were treated with DENV DIPs, delivering 1000 molecules of DI290 RNA per cell or with DIP storage buffer as a negative control. The viral titers in the supernatant from control-treated cells were approximately 10^3^ CCID_50_/mL. However, viral titers in DENV DIP-treated MDM, THP-1 and HFF-1 cells were either at the limit of detection or below (Fig. 1E, 2D and Supplementary Fig. 1C), indicating that DENV DIPs effectively inhibit DENV-2 replication *in vitro* in macrophages and fibroblasts, consistent with previous observations in other cell types.^5,6^ In summary, these findings demonstrate that DENV DIPs stimulate cellular innate immune responses with potent antiviral activity in human MDM, THP-1 and fibroblast cells.

### LNPs delivering DI290 RNA inhibit DENV-2 *in vitro* and *in vivo*

LNPs delivering DI290 RNA were used to treat MDM and HFF-1 cells *in vitro* to validate the stimulation of interferon responses,^28^ as observed with DIP treatment. MDMs and HFF-1 cells were treated with LNP-DI290 delivering 250 ng of DI290 for 24 hours. RT-qPCR analysis for RNA from treated and untreated cells showed that LNP-DI290 significantly upregulated IFN-β, IFN-λ, RIG-I and ISG15 mRNA levels in MDMs and HFF-1 cells compared to cells treated with empty LNPs and untreated cells after 24 hours (Fig. 3A-D and 4A-C). These results demonstrate that LNPs effectively deliver DI290 RNA, stimulating innate immune responses and upregulating IFNs and ISGs in MDMs and HFF-1 cells.

**Figure 3.**
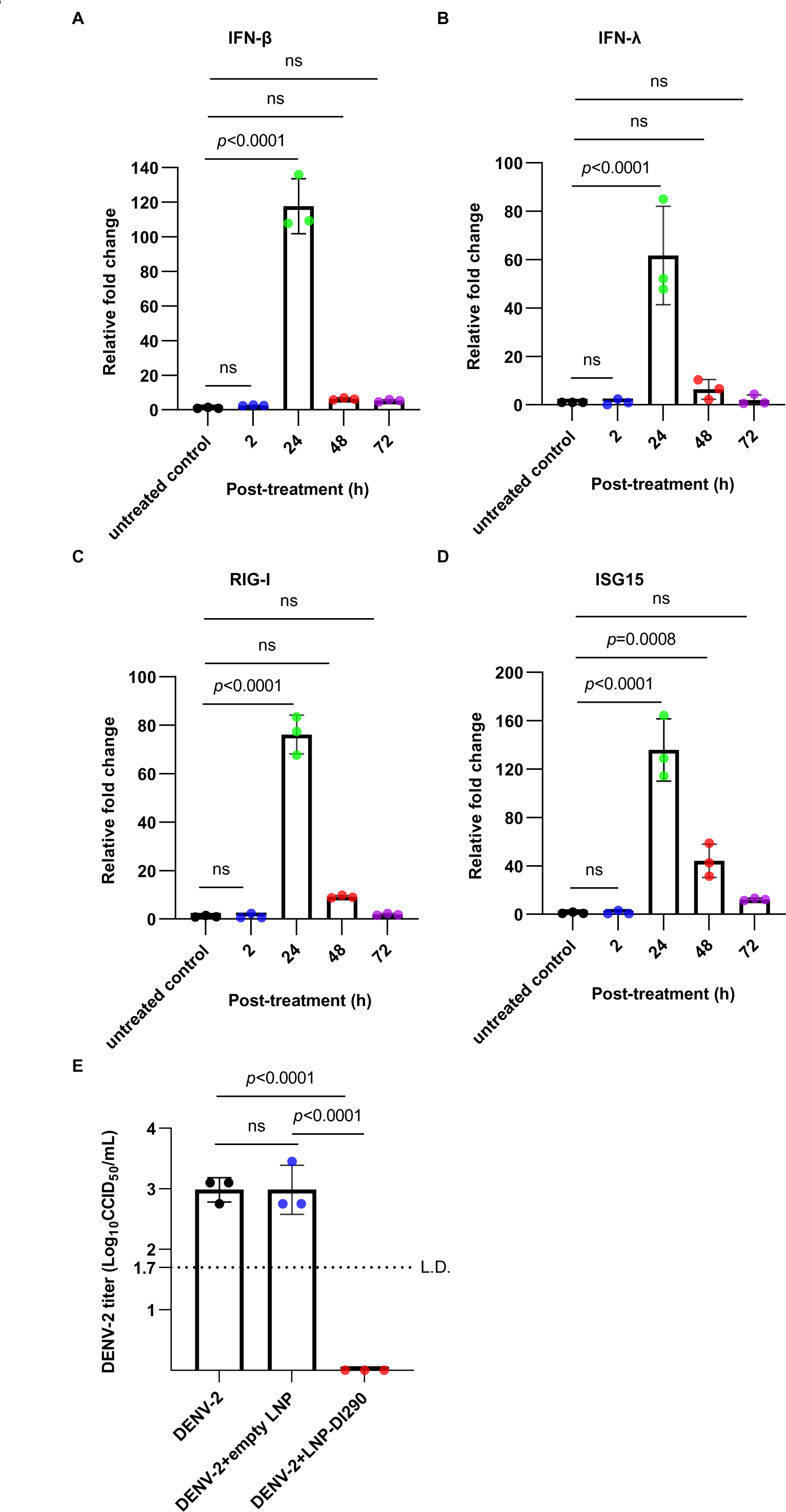
LNP-DI290 stimulates host innate immune responses in human primary MDMs and inhibits DENV-2 replication. (A-D) MDMs were treated with LNP-DI290 (at a dosage equivalent to 25 pg DI290 RNA per cell) for 2, 24, 48 and 72 h. Total RNA was extracted from the cells and the levels of IFN-β, IFN-λ, RIG-I and ISG15 mRNA were quantified by RT-qPCR. The fold change relative to the untreated control cells was calculated (n=3). (E) MDM were infected with DENV-2 (MOI=1 CCID_50_ per cell). After 3 h, the cells were washed with 1X PBS and incubated with either culture medium alone, medium containing either empty LNP or LNP-DI290 (at a dosage equivalent to 25 pg DI290 RNA per cell). DENV-2 titers in culture supernatant were measured by CCID_50_ assay at 3 days post-infection (n=3). Data are shown as the mean ± SD. Statistical analysis was performed by one-way ANOVA. L.D.: limit of detection. ns: not significant.

**Figure 4.**
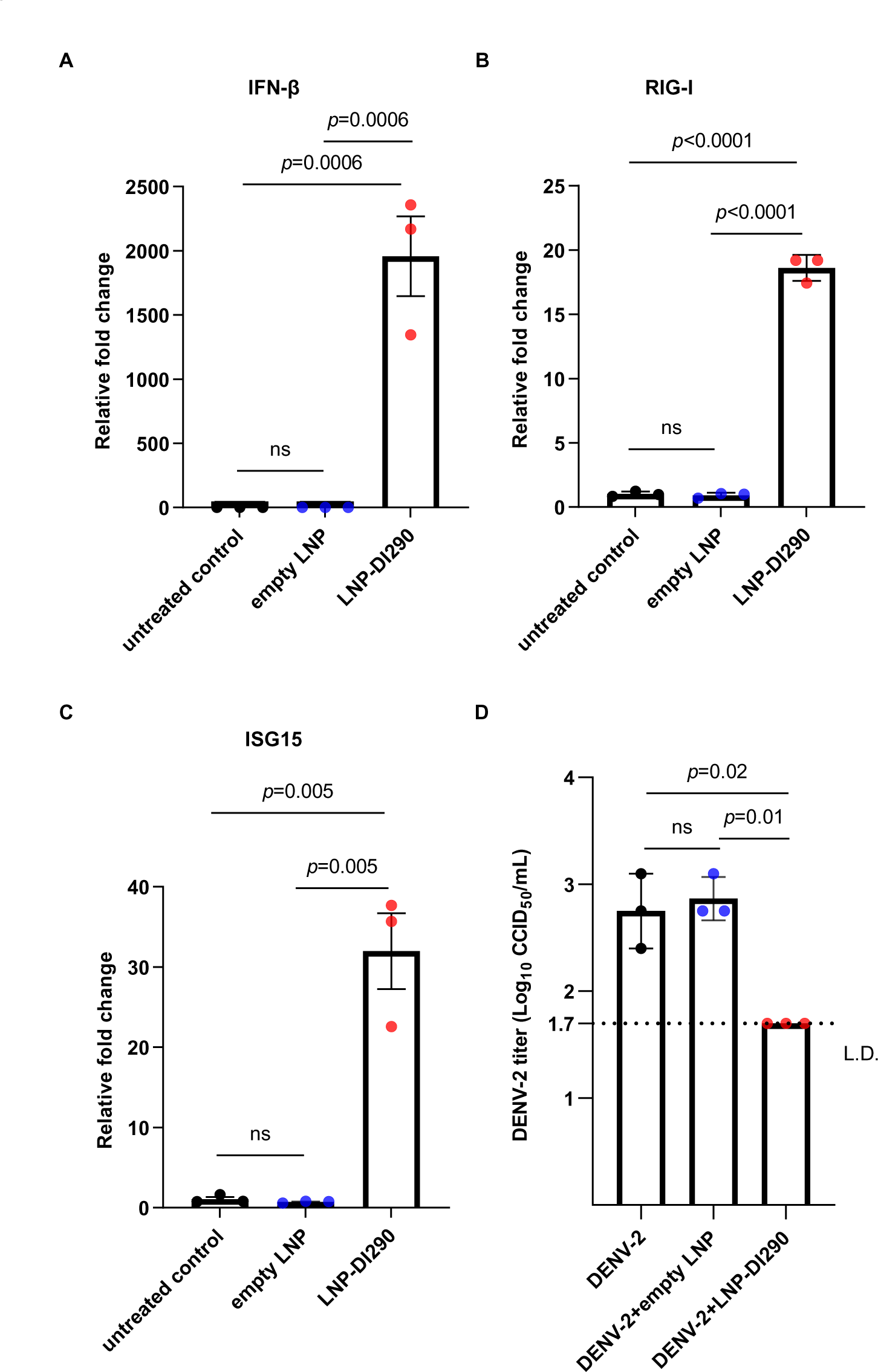
LNP-DI290 stimulates host innate immune responses in HFF-1 cells and inhibits DENV-2 replication. (A-C) HFF-1 cells were treated with empty LNP or LNP-DI290 (at a dosage equivalent to 25 pg DI290 RNA per cell) for 24 h. Total RNA was extracted from the cells and the levels of IFN-β, RIG-I and ISG15 mRNA were quantified by RT-qPCR. The fold change relative to the untreated control cells was calculated (n=3). (D) HFF-1 cells were infected with DENV-2 (MOI=0.1 CCID_50_ per cell). After 3 h, the cells were washed with 1×PBS and incubated with either culture medium alone, medium containing empty LNP or LNP-DI290 (at a dosage equivalent to 25 pg DI290 RNA per cell). DENV-2 titers in culture supernatant were measured by CCID_50_ assay at 3 days post-infection (n=3). Data are shown as the mean ± SD. Statistical analysis was performed by one-way ANOVA. L.D.: limit of detection. ns: not significant.

Subsequently, MDMs and HFF-1 cells were infected with DENV-2 and treated with LNP-290 as in previous experiments. As depicted in figures 3E and 4D, cells treated with LNP-290 exhibited strong inhibition of DENV-2 compared to cells treated with empty LNPs and untreated cells.

LNPs have the capability to deliver significant amounts of RNA *in vivo*, which prompted us to evaluate the antiviral efficacy of LNP-DI290 in a DENV-mouse model. IFNAR-deficient mice are genetically modified to have a specific deletion of the gene encoding the interferon-alpha/beta receptor 1 (IFNAR-deficient). This receptor is a critical component of the signaling pathway for type I interferons. By knocking out the *Ifnar1* gene, these mice lack the receptor necessary for cells to respond to type I interferons, thereby exhibiting heightened susceptibility to infection by viruses, including DENV.^29^ IFNAR-deficient mice were infected by intraperitoneal (*i.p.*) injection of mouse-adapted D220 DENV-2 strain (10^4^ CCID_50_ per mouse), followed by treatment with LNP-DI290 delivering 15 µg of RNA, empty LNPs or buffer only at 4 hours post-infection (h.p.i) (Fig. 5A). Serum and spleen samples were collected at 72 h.p.i. and subjected to assays for infectious virus or viral RNA. Infectious D220 was detected in the serum and spleen at >3 Log_10_ CCID_50_/mL, while infectious virus was undetectable in the spleen and serum samples from mice treated with LNP-DI290 (Fig. 5B-C). Conversely, all samples from untreated and empty LNP-treated mice exhibited measurable titers of D220, indicating the antiviral activity of LNP-DI290 *in vivo*.

**Figure 5.**
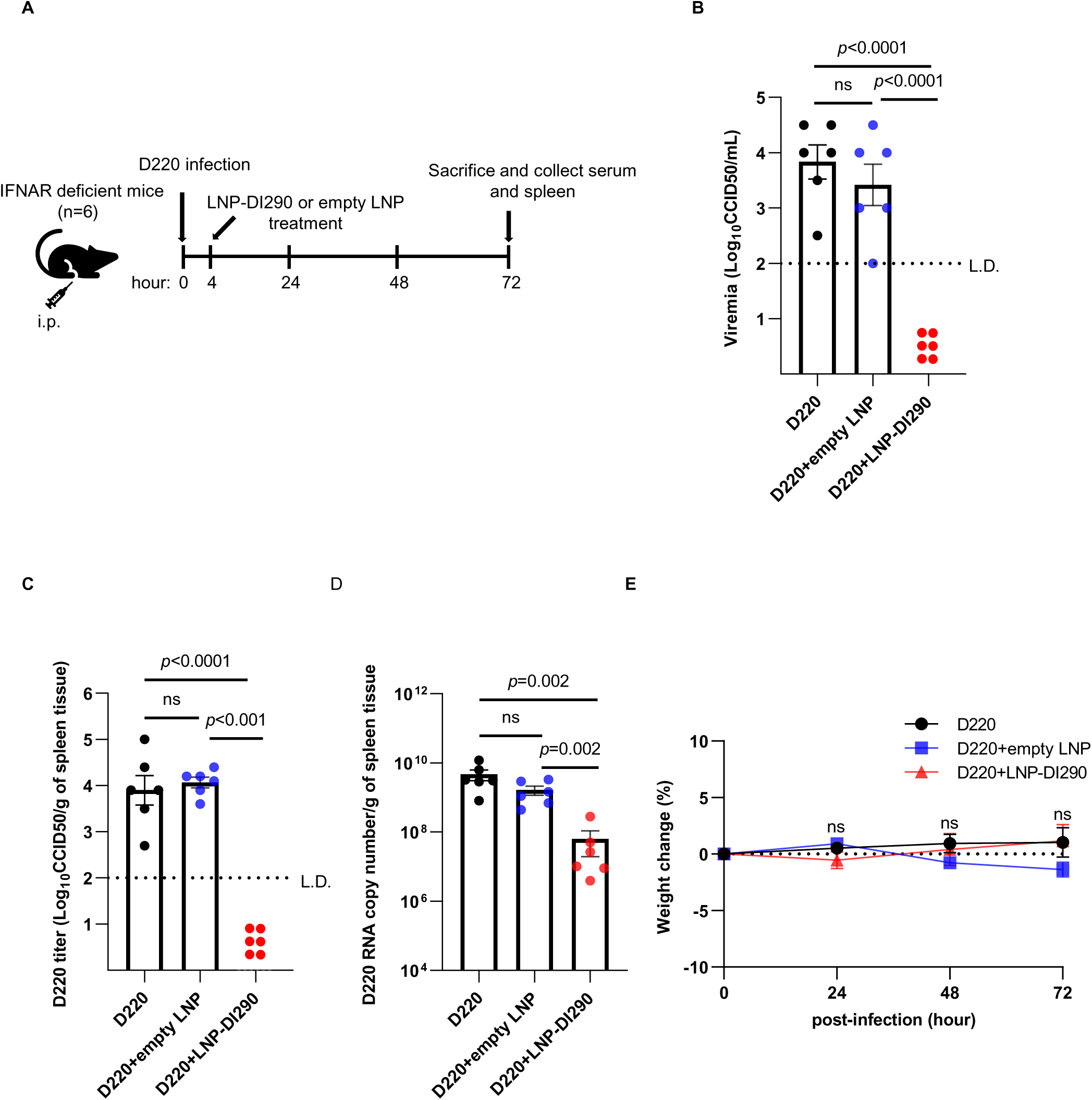
LNP-DI290 inhibits DENV replication in infected IFNAR-deficient mice. (A) Schematic representation of the experimental design. IFNAR-deficient mice were *i.p.* infected with mouse-adapted DENV-2 strain D220 (10^4^ CCID_50_ per mouse). After 4 h infection, the mice were *i.p.* treated with either empty LNP or LNP-DI290 (15 μg DI290 RNA per mouse). (B and C) Viremia and virus titers in spleen were measured by CCID_50_ assay at 72 h post-infection (n=6). (D) The level of DENV-2 D220 RNA in spleen was measured by RT-qPCR using primers for the D220 NS5 region (n=6). (E) The percentage of weight change was monitored at 72 h post-infection (n=6). Data are shown as the mean ± standard error of mean (SEM). Statistical analysis was performed by one-way ANOVA (B-C and E) or Kolmogorov-Smirnov exact test (D). L.D.: limit of detection. ns: not significant.

RT-qPCR analysis of total RNA isolated from splenic samples of all mice showed a significant decrease in viral RNA in LNP-DI290 treated mice compared to mice treated with empty LNPs or buffer (Fig. 5D). This indicates that mice treated with LNP-DI290 were infected by DENV-2 but did not result in measurable levels of viremia *in vivo*. Finally, LNP-DI290 treated mice did not observe significant weight loss, suggesting that DENV infection and LNP-DI290 treatment did not induce an acute metabolic or other adverse change over the course of the experiment. The combined data show that LNP-DI290 delivering 15 µg of DI290 RNA strongly inhibited D220 replication and reduced viremia in IFNAR-deficient mice without apparent adverse effects.

### LNP-DI290 can protect mice from DENV infection independent of IRF3/7

DENV infection leads to the activation of innate immune responses involving several pattern recognition receptors (PRRs) including Toll-like receptor (TLR) 3,^30^ TLR7,^31^ and the RLRs RIG-I and MDA5,^32^ among others. IRF3 and IRF7, the two family members with the greatest structural homology, are principal mediators of type I IFN induction.^reviewed^ ^in^ ^33^ We infected IRF3/IRF7 double knockout (DKO) mice with D220 and then treated them with LNP-DI290, empty LNP or buffer (Fig. 6A). Viremia in serum at >2.5 Log_10_ CCID_50_/mL and viral titre in spleen tissue at >3 Log_10_ CCID_50_/mL was detected after 3 d.p.i (Fig. 6B-C). However, in LNP-DI290 treated mice, no virus was detected in serum or spleen samples. We measured viral RNA in total spleen RNA samples from all mice (Fig. 6D), but no viral RNA was detected in mice treated with LNP-DI290 (Fig. 6D). As observed for IFNAR-deficient mice, significant weight loss was not observed over the course of the experiment in these mice (Fig. 6E). Given the importance of RIG-I and MDA5 in establishing innate immune responses to DENV infection ^32^, this outcome indicates, surprisingly, that DI290 RNA induced innate immune responses in mice strongly inhibited DENV in an IRF3/7 independent manner.

**Figure 6.**
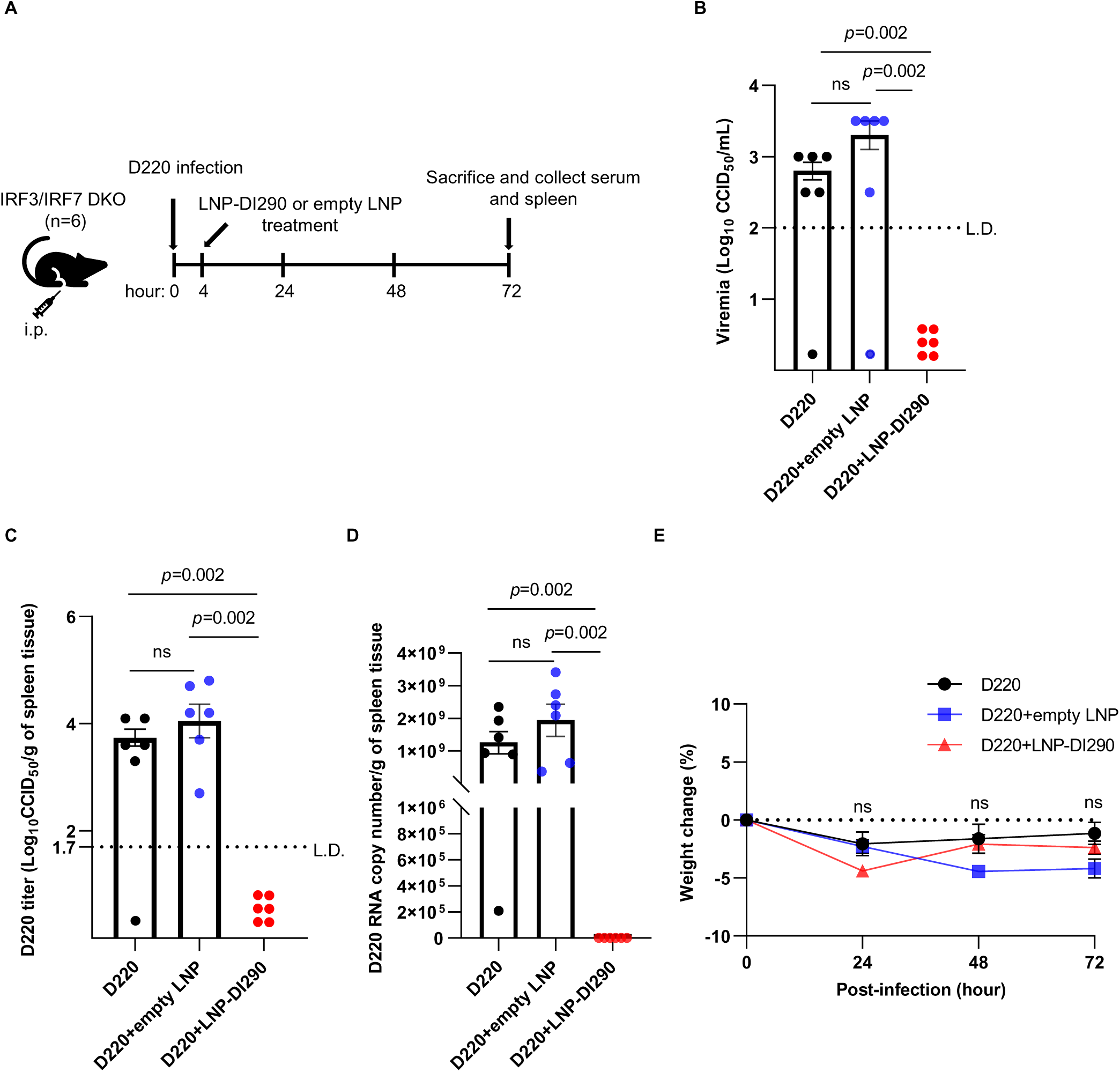
LNP-DI290 block DENV replication in infected IRF3/IRF7 DKO mice. (A) Schematic representation of the experimental design. IRF3/IRF7 DKO mice were *i.p*. infected with D220 (10^4^ CCID_50_ per mouse). After 4 h infection, the mice were *i.p.* treated with either empty LNP or LNP-DI290 (15 μg DI290 RNA per mouse). (B and C) Viremia and virus titres in spleen were measured by CCID_50_ assay at 72 h post-infection (n=6). (D) The level of D220 DENV-2 RNA in spleen was measured by RT-qPCR using primers for the D220 NS5 region (n=6). (E) The percentage of weight change was monitored for at 72 h post-infection (n=6). Data are shown as the mean ± SEM. Kolmogorov-Smirnov exact test was used for statistical analysis. L.D.: limit of detection. ns: not significant.

### Activation of innate immune response including IFN-γ following treatment of DENV DIPs and LNP-DI290 *in vivo* and *in vitro*

Activation of the innate immune response following treatment with DENV DIPs and LNP-DI290 was examined both *in vivo* and *in vitro*. To investigate host responses after treatment with DENV DIPs/LNP-DI290, we employed RNA sequencing (RNA-Seq) to quantitatively measure the expression levels of transcripts in spleen tissues from both C57BL/6J and IFNAR-deficient mice treated with LNP-DI290 or empty LNPs. Additionally, we analyzed RNA from MDMs treated with LNP-DI290 and empty LNPs. We also included DENV DIP-treated MDMs to compare induction patterns following LNP treatment versus DIP treatment.

Approximately 10% of genes in the human genome have the potential to be regulated by IFNs.^34^ The RNA-Seq analysis revealed substantial transcriptional alterations in host cells induced by DENV DIP/LNP-DI290 treatment (Supp. #. RNA Seq raw data). A total of 87 upregulated genes were commonly identified in mouse spleen tissues and human MDMs (Fig. 7). Notably, over 45 shared upregulated genes in response to these treatments represented interferon-stimulated genes (ISGs), including ZPB1, MX1, ISG15, IFIT1-3, IFITM3, and OAS2-3, among others. IFNs, as secreted cytokines, activate signal transduction cascades leading to the induction of hundreds of ISGs. ISGs upregulated in MDMs and mice in this study, which were previously reported to inhibit DENV replication, include ISG15,^35^ 2′,5′-oligoadenylate synthetase (OAS) 3,^36^ IFITM3,^37^ BST2 (Viperin),^37,38^ and CMPK2.^39^ In addition, PRRs, including RIG-I, MDA5 (IFIH1) and LGP2, were upregulated by DI290 RNA even in IFNAR-deficient mice where IFNα/β is impeded.

**Figure 7.**
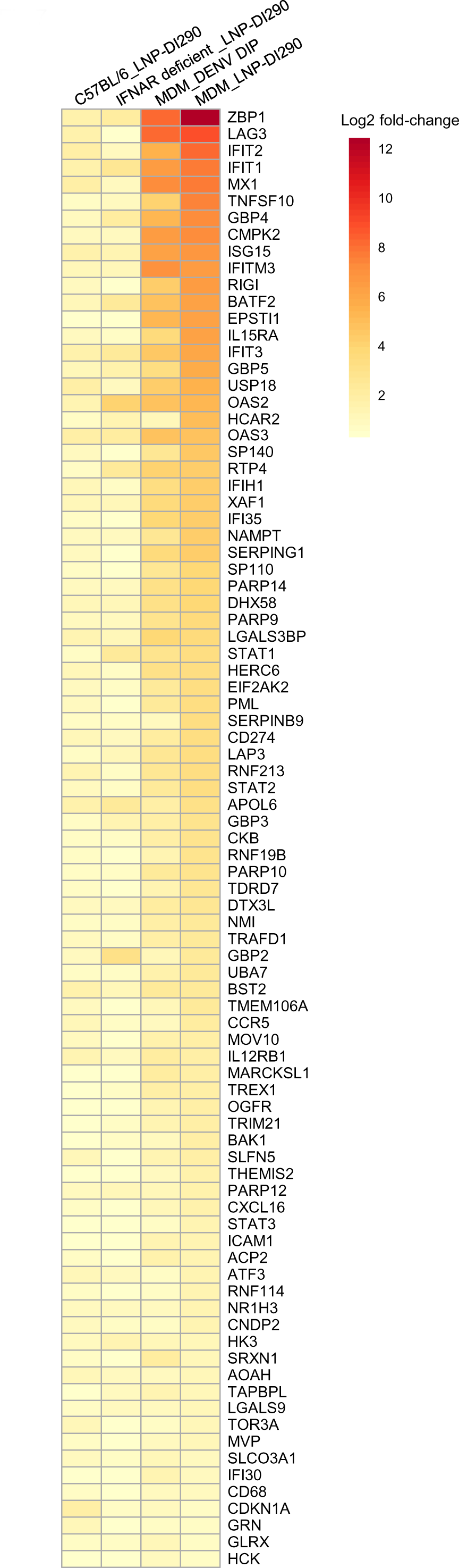
Results of EdgeR analysis. Heatmap shows log_2_ fold-changes of 87 DEGs that were significantly upregulated in all four experimental groups. DEGs are ordered by log_2_ fold-change in the LNP-DI290 treated MDM (MDM_LNP-DI290) group.

To explore the cellular responses further, the outcomes of RNA-Seq analysis were subjected to Ingenuity Pathway Analysis (IPA) Core Analysis to identify common upregulated or downregulated upstream regulators (USRs) after DIP or LNP-290 treatment. In figure 8, the heatmaps show activation z-scores of cytokine/chemokine USRs (Fig. 8A) and transcriptional regulators USRs (Fig 8B) that were significantly upregulated. The analysis showed activation of 117 cytokines/chemokines that include IFN-γ (IFNG), TNF, IFN-α (IFNA1 and IFNA2), IFN-β (IFNB1) and IFN-λ (IFNL1). The activation z-score of IL-27 is consistent with the regulation of both innate and adaptive immunity USRs.^40^ IL-27 can induce IFN-γ and inflammatory mediators from T lymphocytes and innate immune cells.^41^ The activation z-scores of IRFs included IRF1, IRF3, IRF5, IRF7 and IRF9 (Fig 8B). Other activation z-scores for transcription factors include Zbtb10, which is involved in dendritic cell activation,^42^ STAT1, STAT2, and NFκB (NFKB1, NFKB2), which regulate inflammatory responses. Taken together, the observed modulation of key inflammatory regulators, particularly the activation of IFNs and associated immune effectors, demonstrates that DENV DIP/LNP-DI290 treatment triggers a robust innate immune response when stimulated by DI290 RNA *in vitro* and *in vivo*.

**Figure 8:**
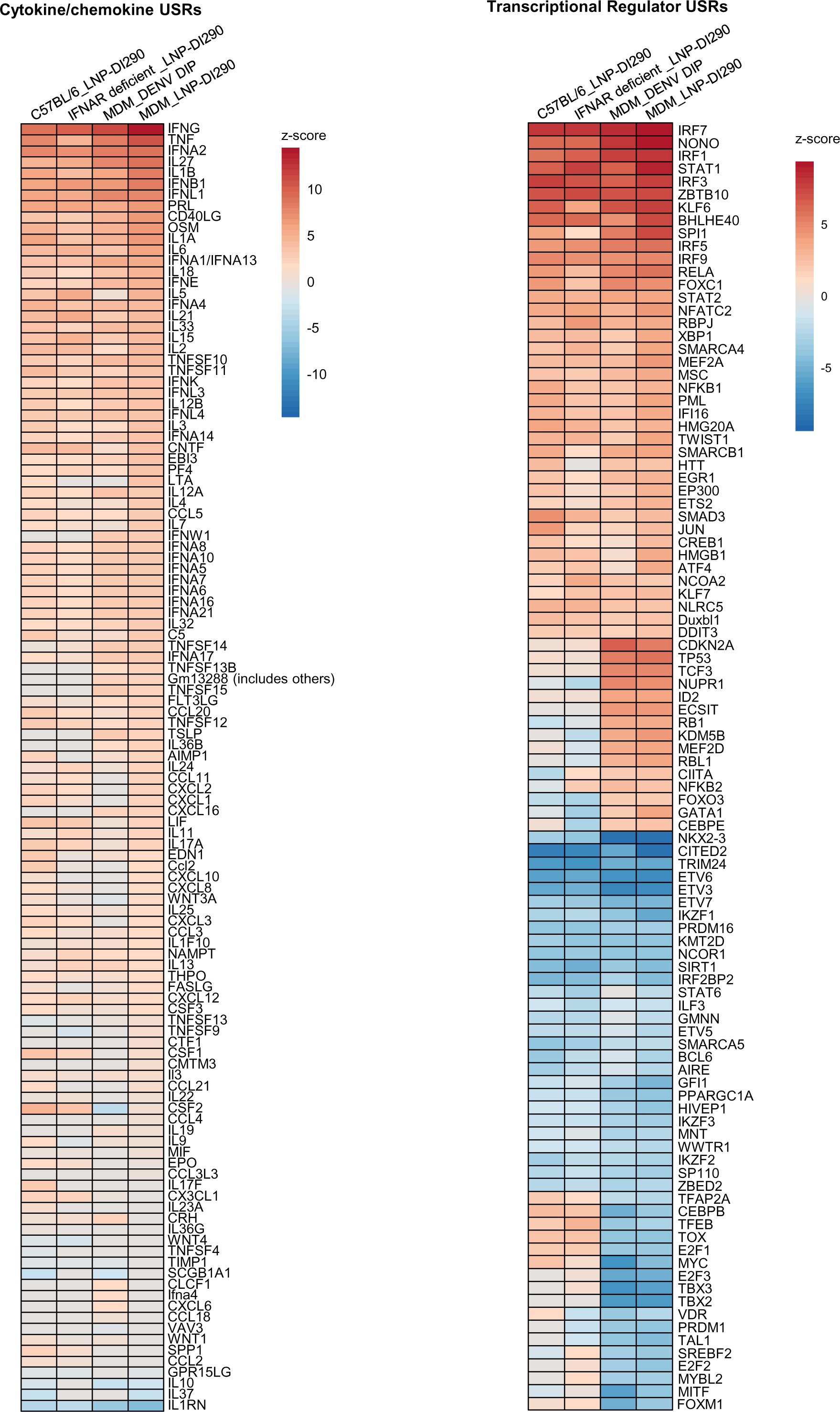
Results of IPA Core Analysis. (A) Heatmap shows activation z-scores of 117 cytokine/chemokine USRs that displayed significant activation or inhibition in at least one experimental group. Any USR that was not significant in a particular group was given a z-score of zero. USRs are ordered by z-score in the MDM group treated with LNP290 (MDM_LNP-DI290). (B) Heatmap shows activation z-scores of transcription factor USRs that displayed significant activation or inhibition in at least one experimental group. The 100 most activated/inhibited USRs across all groups are shown. Any USR that was not significant in a particular group was given a z-score of zero. USRs are ordered by z-score in the MDM_LNP-DI290 group.

## Discussion

Here, we showed that DI290 RNA, a naturally derived DENV-2 DI RNA [4], triggers host cell antiviral responses, effectively suppressing DENV infection both *in vitro* and *in vivo*. DI RNA belongs to the class of defective viral genomes (DVGs), which emerge during viral replication within cells and are packaged and released as non-infectious virus-like particles (VLPs) alongside infectious wild-type viruses. ^10,43^. The presence of these VLPs in a virus stock is linked with reduced viral replication and virulence, indicating attenuation.^14,44–46^ In this study, we utilized DI290 RNA delivered through DIPs or LNPs to activate cellular innate immune responses capable of counteracting DENV replication. DENV employs well-characterized mechanisms to evade the innate antiviral response. For instance, DENV NS3 can inhibit RIG-I activation,^47^ while the DENV NS5 protein disrupts IFN signaling by binding to and degrading STAT2.^48–50^ For Sendai virus, DVGs possess the capability to overcome virus-mediated inhibition of IFN induction, potentially through preferential binding of a DVG to RIG-I or MDA5 compared to the wild-type genome.^14^ It is plausible that DI290 may exhibit preferential binding to an RLR such as RIG-I, thereby inducing IFN responses in DENV-infected cells. This possibility could be further investigated in future studies, for instance, utilizing techniques like RNAscope.^51^

The IPA analysis of RNAseq data revealed that DI RNA notably upregulated IFN-γ signaling in MDM cells, IFNAR-deficient mice and C57BL/6J mice. IFNAR-deficient mice are susceptible to DENV infection due to impaired IFN-α/β signaling and diminished antiviral responses. However, treatment with LNP-DI290 significantly reduced viremia and virus replication in splenic tissue. Our analysis suggests that IFN-γ signaling may have facilitated the upregulation of ISGs, inhibiting DENV replication in IFNAR mice. Notably, IFN-γ signaling is protective against DENV pathogenesis in vaccinated individuals, ^52^ and in IFNAR mice.^53^ IPA z-scores indicated a significant increase in TNF, IFN-γ, and IFN-λ1. This finding supports the notion that DI290 RNA exerts its effect in IFNAR-deficient mice by upregulating antiviral activity. Further support for this possibility was observed in experiments using IRF3/IRF7 DKO mice. In all systems treated with LNP-DI290 or DIPs, similar patterns of pathway activation in DEGs, cytokine/chemokine and transcriptional regulators were evident. The PRRs that trigger antiviral responses upon infection, including TLR3, TLR7, RIG-I and MDA5, induce both type I and type III IFNs via IRF3 and IRF7. Notably, viral infection was not detected in IRF3/IRF7 DKO mice treated with LNP-DI290, which lack IFN type I expression but retain functional IFN type I receptors. DI290 RNA may stimulate “noncanonical” signaling pathways, ^54^ in MDMs, IFNAR mice and IRF3/IRF7 DKO mice, resulting in the upregulation of IFN-driven antiviral responses.

Activation of signaling pathways by DI RNA may lead to unintended consequences, including off-target effects that could be harmful. Excessive activation of the immune system, particularly if sustained, can result in immunotoxicity. We conducted preliminary testing of LNP-DI290 in IFNAR-deficient and C57BL/6J mice, where the delivery of DI290 RNA in IFNAR-deficient and IRF3/IRF7 DKO mice did not cause any significant levels of weight loss, an indicator no serious metabolic change. It is possible that the efficacy and safety of DI290 may depend on the dose, and finding the right balance between stimulating the immune response and avoiding adverse effects will be crucial.

This study showed that LNP-DI290 administered to infected mice 4 h.p.i. prevented viremia. Treatment timing in this model was limited due to rapid peak viremia (at 3 d.p.i.) and viral clearance (at 4-5 d.p.i) in *Ifnar^-/-^* mice. Future studies to investigate if prophylactic LNP-DI290 application is protective, or if administration at a later time point post-infection provides protection from infection, is warranted but a different approach and animal model may be necessary. For example, AG129 mice, which have KO for both type I and type II IFN receptors, permit higher levels of viral replication compared to IFNAR mice. A recent study showed that both models resulted in a similar time course with respect to viremia. Interestingly, aspects of severe dengue disease were recapitulated by both models involving DENV-mediated inflammation and tissue damage in the gastrointestinal tract (GIT).^55^ The GIT pathology occurred late in the disease after the peak of viremia. Infected macrophage but not endothelial cells in the GIT were reported. Whether DI290 RNA therapy ameliorates or exacerbates GIT pathology could help define the utility of DI290 therapy during the late stages of DENV disease.

In summary, we have demonstrated that DI290 RNA delivered by DIPs or LNP-290 activates innate immune responses in macrophages, a primary target for DENV infection, and strongly inhibits DENV infection in THP-1 and human MDMs. LNP-DI290 strongly inhibits DENV replication in IFNAR-deficient mice and IRF3/IRF7 DKO mice. Control of DENV replication was associated with solid upregulation of IFN signaling pathways. Under the conditions tested, a single dose post-infection treatment did not appear to cause adverse events in these animal models, indicating that DI290 RNA has potential as a therapy for DENV infection.

## Materials and Methods

### Ethics statement

All mouse work was conducted in accordance with the “Australian code for the care and use of animals for scientific purposes” as defined by the National Health and Medical Research Council of Australia. All animal procedures were conducted in a dedicated suite in a biosafety level-3 (PC3) facility at the QIMR Berghofer Medical Research Institute (Australian Department of Agriculture, Water and the Environment certification Q2326 and Office of the Gene Technology Regulator certification 3445) as previously described ^56^. All work was approved by the QIMR Berghofer Medical Research Institute Animal Ethics Committee (P2277, A2002-600). Mice were euthanized using carbon dioxide asphyxiation. Overt clinical signs of mice were scored as described in ^57^ on a scale of 0–3 (Diseases scores) according to posture, activity and fur ruffling, with a score of 0 meaning no clinical signs were observed.

Breeding and use of GM mice were approved under a Notifiable Low Risk Dealing (NLRD) Identifier: NLRD_Harrich_Aug_2023: NLRD 1.1(a), NLRD 2.1(d), NLRD 2.1(j), NLRD 2.1(l).

### Cell culture

Vero E6 (ATCC CRL-1586) and HFF-1 (ATCC SCRC-1401) cells were maintained in Dulbecco’s Modified Eagle medium (DMEM) (Thermo Fisher) supplemented with 10% (v/v) fetal bovine serum (FBS) (Thermo Fisher) and 1% (v/v) penicillin-streptomycin (Thermo Fisher). DENV DIP-producing cell line, HEK-DI-290-ORF, was cultured in HEK GM Serum-Free Medium (Sartorius) supplemented with 1% (v/v) penicillin-streptomycin, 1% (v/v) GlutaMAX-I (Thermo Fisher) and 50 nM PEP005 (Adooq Bioscience) during DENV DIP production ^6^. The human monocytic cell line THP-1 (a gift from Prof. Andreas Suhrbier, QIMR Berghofer Medical Research Institute) was maintained in RPMI 1640 medium supplemented with 10% (v/v) FBS and 1% (v/v) penicillin-streptomycin. THP-1 cells were differentiated into macrophage-like cells by stimulation of 50 ng/mL phorbol myristate acetate (PMA) for 48 h. The THP-1 cells were then cultured for 24 h with PMA-containing medium replaced with a complete medium. All cells were incubated at 37 °C in a humidified 5% CO_2_ atmosphere. C6/36 cells (a gift from Prof. Andreas Suhrbier, QIMR Berghofer Medical Research Institute) were grown in DMEM supplemented with 10% (v/v) FBS and 1% (v/v) penicillin-streptomycin. Cells were checked for mycoplasma using MycoAlert Mycoplasma Detection Kit (Lonza Bioscience). Cell lines are routinely authenticated in-house by Short Tandem Repeat profiling.

### Isolation of monocytes from peripheral blood mononuclear cells (PBMC)

PBMCs were isolated from a healthy donor’s buffy coat supplied by the Australian Red Cross Blood Service using Ficoll density gradient centrifugation as previously described.^58^ After PBMC isolation, cells were washed and resuspended in PBS containing 2% FBS and 1 mM EDTA. Monocytes were isolated from PBMCs using EasySep™ Human CD14 Positive Selection Kit according to the manufacturer’s instructions (Stemcell Technologies).

### Differentiation of human monocyte to MDMs

MDMs were differentiated as previously described.^59^ Briefly, Isolated monocytes were cultured in RPMI 1640 (Thermo Fisher) supplemented with 10% (v/v) FBS, 1% (v/v) penicillin-streptomycin, 10 ng/mL macrophage colony-stimulating factor (M-CSF) (Thermo Fisher) and 1 ng/mL granulocyte-macrophage colony-stimulating factor (GM-CSF) (Thermo Fisher) for 5 days. Medium was changed every 2 days.

### DENV DIP production, precipitation and chromatographic purification

HEK-DI-290-ORF cells were seeded at 2.5 × 10^5^ cells/ml and grown in a ProCulture glass spinner flask (Corning) using a magnetic stir plate (Thermo Fisher) set at 65 rpm. After 48 h, the culture supernatant containing DENV DIPs was collected, filtered through a 0.22 μm filter (Merck Millipore), and the clarified supernatant was stored at 4 °C. The clarified supernatant was processed through the tangential flow filtration (TFF) system (AKTA Flux S, Cytiva), which was equipped with 100 kDa molecular weight cut-off filter (UFP-100-C-H24LA hollow fiber cartridge) and equilibrated with sterile 1X PBS. Upon equilibration, the clarified supernatant was added to the TFF chamber and flowed through 54 mL/min and a permeate pump rate of 5 mL/min. The transmembrane pressure was maintained below 0.5 bar. The resulting concentrated retentate was then filtered through a 0.22 μm filter and added with stabilizing buffer containing 2% (v/v) gelatin hydrolysate (Merck G0262) and 5% (v/v) sorbitol (Merck S1876) before CHT ceramic hydroxyapatite (Bio-Rad Laboratories) chromatographic purification of DENV DIPs.

CHT chromatography column purification was performed as previously described ^6^. The resulting purified DENV DIPs were then concentrated using an Amicon Ultra-15 centrifugal filter unit (50 kDa cutoff) until the volume was reduced to around 1-1.5 mL. Quantification of DI290 RNA copies was performed by qPCR analysis.

### LNP-DI290 Formulation

LNPs encapsulating DI290 RNA, were created using the hydration method.^60^ Briefly, required amounts of lipid and PEG2000-C16Ceramide were mixed with DI290 RNA at an N:P ratio of 4 in a sucrose-containing water/tert-butanol (1:1 v/v) co-solvent system. DOTAP, cholesterol, DOPE and PEG2000-C16Ceramide with a molar ratio of 50:35:5:10 was used. The mixture was then snap-frozen and freeze-dried overnight. Freeze-dried matrix was then hydrated with sterile water immediately before use at a concentration of 0.1mg/ml D1290 final.

### Analysis of DENV DIP and LNP-DI290 antiviral activity in vitro

HFF-1, MDM and THP-1 cells were seeded in a 48-well plates with 10,000 cells per well. THP-1 cells were induced for differentiation, as described above. The next day, the cells were infected with DENV-2 (MOI=0.1 CCID_50_ per cell for THP-1 and MOI=1 CCID_50_ per cell for THP-1 and MDM). After 3 h, the cells were washed with 1X PBS and incubated with either culture medium alone, medium containing DENV DIP (at a dosage equivalent to 1,000 DI290 RNA copies (=0.2 fg) per cell), empty LNP or LNP-DI290 (at a dosage equivalent to 1.6 × 10^8^ DI290 RNA copies (=25 pg) per cell). At 3 days post-infection, the viral titers were determined by CCID_50_ assays as previously described ^6,61^.

### Gene expression analysis

HFF-1, MDM and THP-1 cells were seeded at a density of 50,000 each well in a 48-well plate. THP-1 cells were induced for differentiation, as described above. The next day, the cells were treated with culture medium only or medium containing either DENV DIP (equivalent to 1,000 DI290 RNA copies (= 0.2 fg) / cell), empty LNP or LNP-DI290 (equivalent to 1.6 × 108 DI290 RNA copies (= 25 pg) / cell). Total RNA from cells was extracted after 2 h, 24 h, 48 h and 72 h treatment using RNeasy Kit (Qiagen) and mRNA expression levels of Ribosomal Protein L13a (RPL13A), IFN-β, RIG-I and ISG15 were determined using Luna Universal One-Step RT-qPCR Kit according to the manufacturer’s instructions (New England Biolabs). The primer sequences are available upon request. The data was normalized to RPL13A as the reference gene ^62^ and presented as fold change relative to untreated control cells using ΔΔC_t_ method.

### Mouse infection

All mice were bred in-house and housed at QIMR Berghofer Medical Research Institute, Brisbane, QLD, Australia. IFNAR-deficient mice,^63^ and IRF3/IRF7 DKO mice,^64^ were infected *i.p.* with 1 × 10^4^ CCID_50_ of DENV-2 strain D220,^65^ for 4 h. Mock-infected mice were injected i.p. with PBS. After 4 h infection, the mice were treated i.p. with either LNP-DI290 containing 15 μg of DI290 RNA or empty LNPs.

Following treatment, body weight and general conditions were monitored daily for up to 3 days. The mice were euthanized using CO_2_ at 3 days post infection. Blood was harvested through heart puncture and the spleen was collected and homogenized in 1 ml RPMI 1640 containing 2% FBS. The homogenate (200 μl) was mixed with Trizol (Thermo Fisher) for RNA extraction, followed by quantification of viral RNA using RT-qPCR. Virus titration in both blood and spleen samples was determined by CCID_50_ assays as previously described.^6,61^

### Quantification of viral RNA from spleen samples

After Trizol extraction, the RNA pellet was resuspended in 50 µl of RNase-free H_2_O and 1 µl of the resuspended samples was used for quantification of the DENV-2 D220 NS5 RNA regions using Luna Universal One-Step RT-qPCR Kit (New England Biolabs) in a 10-ul reaction volume. The thermal cycling conditions were as follows: 15 min at 55 °C, 2min at 95 °C and 40 cycles of 5 sec at 95 °C, 45 sec at 55 °C (data acquisition) and 15 sec at 72 °C. The primer sequences are available on request.

### RNA Seq sample library preparation and analysis

For the preparation of MDM mRNA, MDMs were seed in 48-well plates with 30,000 cells per well and treated with either culture medium alone, medium containing DENV DIP (at a dosage equivalent to 0.2 fg DI290 RNA copies per cell, n=3), empty LNP (n=3) or LNP-DI290 (at a dosage equivalent to 25 pg DI290 RNA copies per cell, n=3). After 24 treatments, total RNA was extracted from the cells using RNeasy Kit (Qiagen) according to the manufacturer’s instructions. For mRNA preparation from mouse spleens, C57BL/6J (n=6) and IFNAR-deficient (n=5) mice were *i.p.* treated with empty LNP or LNP-DI290 (at a dosage equivalent to 15 μg DI290 RNA per mouse). After 24 hours of treatment, whole spleens were harvested, preserved in RNA*later*^TM^ solution (Thermo Fisher) and homogenized in Trizol (Thermo Fisher). Total spleen RNA was then extracted following the manufacturer’s instructions.

RNA concentration and quality were measured using TapeStation D1kTapeScreen assay (Agilent). cDNA libraries were generated using Illumina TruSeq Stranded mRNA library prep kit and were sequenced using an Illumina Nextseq 2000 Sequencing System to produce 75nt paired-end reads. Sequence reads were aligned to either the GRCm39 v33 mouse or the GRCh38 v44 human reference genomes obtained from Gencode, using STAR aligner v2.7.10. Aligned read counts were calculated for each annotated gene using RSEM v1.3.1, and differential expression was estimated with EdgeR v3.42.4 using the statistical software, R v4.2.0. Only genes with at least 0.5 counts-per-million (approximately 10 aligned reads) were included in the test for differential expression. Significantly differentially expressed genes (DEGs) were analyzed using Ingenuity Pathway Analysis v107193442 (QIAGEN). IPA accepts DEG sets of between 200-3000 genes. Therefore, false discovery rate (FDR) cut-offs were adjusted to keep DEG sets within this range and roughly comparable between experimental groups. q<1×10-2 was used for the C57BL/6J group; q<1×10-5 was used for the IFNAR-deficient group; q<1×10-3 was used for the DIP-treated MDM group; q<1×10-5 was used for the LNP-DI290-treated MDM group.

### Statistical analysis

Statistical analysis was performed using IMB SPSS Statistics (IMB Corp.) and GraphPad Prism (GraphPad software). When the difference in variance was >4, skewness was <-2 or kurtosis was >2, the data were considered non-parametric and the Kolmogorov–Smirnov exact test was performed. Otherwise, the one-way ANOVA or student’s *t*-test were used. *p-*value <0.05 was considered significant.

## Data and code availability

All data generated or analyzed during this study are included in this published article and its supplementary data files and information. Any remaining information can be obtained from the corresponding author upon reasonable request.

## Acknowledgements

This research was generously supported through funding from the Wellcome Trust Innovator Award 219588Z/19/Z to D.H. and the DARPA INTERfering and Co-Evolving Prevention and Therapy (INTERCEPT) program to D.H. and John Aaskov. We sincerely thank John Aaskov for the introduction to dengue virus defective interfering particles field. We also thank Dr. I Anraku for his assistance in managing the PC3 (BSL3) facility at QIMR Berghofer Medical Research Institute and animal house staff from QIMR Berghofer Medical Research Institute for mouse breeding and agistment.

## Author Contributions

M-H.L. and P.M. conducted the experimental work. L.W-E. and Y.T. synthesized nanoportciles. C.B. conducted RNA-seq analysis. B.T., D.L. and L.L. assisted M-H.L. and P.M. throughout the project. Author D.H., A.S., N.A.J.M. and M-H.L. conceived, designed and directed the study. D.H., M-H.L. and P.M. made the figures. D.H., M-H.L. and P.M. analyzed the outcomes and wrote the manuscript. All authors read and approved the manuscript.

