## Supplemental Figure 1 for "Harnessing defective interfering particles and lipid nanoparticles for effective delivery of an anti-dengue virus RNA therapy"

Sup.1

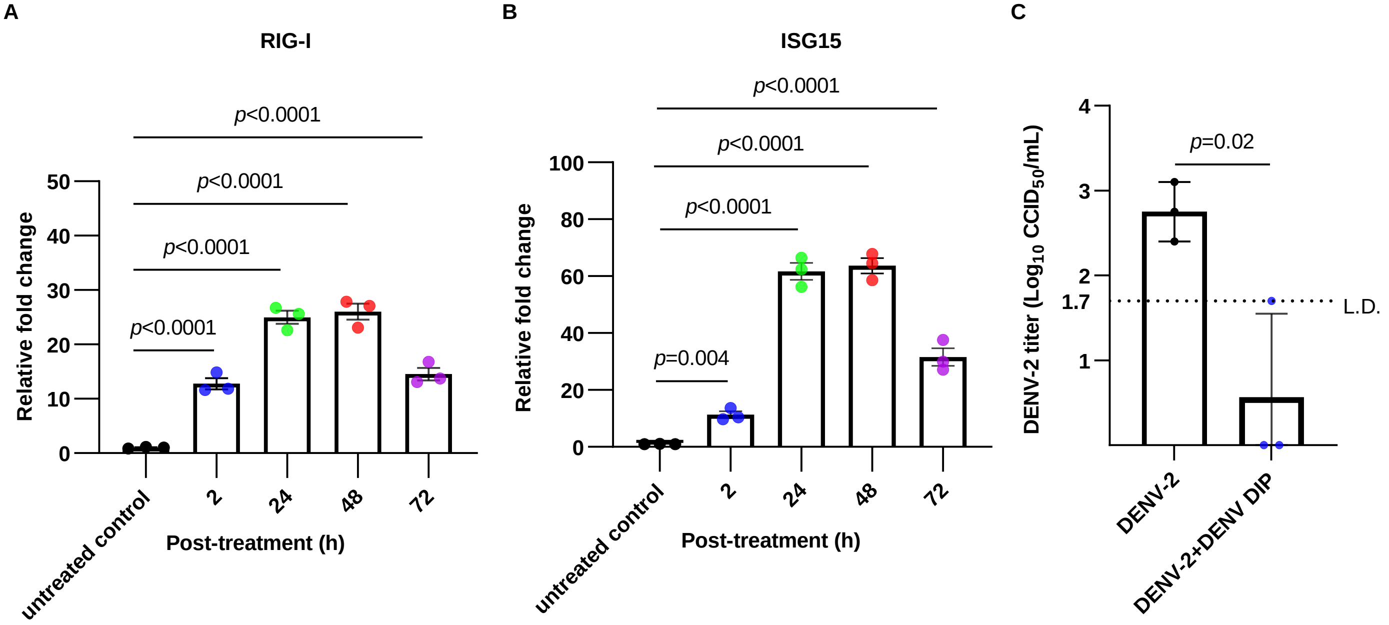


Sup.1. (A-B) THP-1 cells were then treated with DENV DIPs (at a dosage equivalent to 1,000 DI290 RNA copies (=0.2 fg) per cell) for 2, 24, 48 and 72 h. Total RNA was extracted from the cells and the levels of RIG-I and ISG15 mRNA were quantified by RT-qPCR. The fold change relative to the untreated control cells was calculated (n=3). (C) THP-1 cells were infected with DENV-2 (MOI=1 $CCID_{50}$ per cell). After 3 h, the cells were washed with 1X PBS and incubated with culture medium containing DENV DIP (at a dosage equivalent to 1,000 DI290 RNA copies (=0.2 fg) per cell). DENV-2 titers in culture supernatant were measured by $CCID_{50}$ assay at 3 days post-infection (n=3). The data are shown as the mean ± SD. P values were calculated using a two-tailed Student's t test. L.D.: limit of detection.
